# Predicting cell types with supervised contrastive learning on cells and their types

**DOI:** 10.1101/2023.08.08.552379

**Authors:** Yusri Dwi Heryanto, Yao-zhong Zhang, Seiya Imoto

**Affiliations:** Institute of Medical science, the University of Tokyo, Tokyo, 108-0071, Japan

## Abstract

Single-cell RNA-sequencing (scRNA-seq) is a powerful technique that provides high-resolution expression profiling of individual cells. It significantly advances our understanding of cellular diversity and function. Despite its potential, the analysis of scRNA-seq data poses considerable challenges relate to multicollinearity, data imbalance, and batch effect. One of the pivotal tasks in single-cell data analysis is cell type annotation, which classifies cells into discrete types based on their gene expression profiles. In this work, we propose a novel modeling formalism for cell type annotation with a supervised contrastive learning method, named SCLSC (**S**upervised **C**ontrastive **L**earning for **S**ingle **C**ell). Different from the previous usage of contrastive learning in single cell data analysis, we employed the contrastive learning for instance-type pairs instead of instance-instance pairs. More specifically, in the cell type annotation task, the contrastive learning is applied to learn cell and cell type representation that render cells of the same type to be clustered in the new embedding space. Through this approach, the knowledge derived from annotated cells is transferred to the feature representation for scRNA-seq data. The whole training process becomes more efficient when conducting contrastive learning for cell and their types. Our experiment results demonstrate that the proposed SCLSC method consistently achieves superior accuracy in predicting cell types compared to four state-of-the-art methods. SCLSC also performs well in identifying cell types in different batch groups. The simplicity of our method allows for scalability, making it suitable for analyzing datasets with a large number of cells. In a real-world application of SCLSC to monitor the dynamics of immune cell subpopulations over time, SCLSC demonstrates a capability to discriminate cell subtypes of CD19+ B cells that were not present in the training dataset.

## Introduction

In recent years, single-cell RNA sequencing (scRNA-seq) has made remarkable advancements, enabling the analysis of gene expression profiles at the individual cell level. This technology has been instrumental in identifying rare cell populations, defining cell types, and uncovering novel cell states^1,2^. Experts in the field have collected numerous datasets using scRNA-seq, including large-scale initiatives like the Human Cell Atlas^3^ and Tabula Muris Atlas^4^.

The initial step in analyzing single-cell data typically involves annotating cells based on their known and novel cell types. One commonly used strategy, known as “cluster-then-annotate,” involves grouping cells into clusters based on the similarity of their gene expression profiles and manually characterizing them using previously identified cell type markers^4,5^. However, this manual strategy presents several challenges. Firstly, the process of cell type annotation is labor-intensive, requiring extensive literature review of genes specific to each cluster^6^. Secondly, any changes made to the analysis, such as incorporating additional data or adjusting parameters, require the manual reevaluation of all previous annotations. Thirdly, the incompleteness of the current knowledge and researcher subjectivity may contributes to cells mislabeling^7,8^. Lastly, transferring annotations between independent datasets generated by different research groups studying related tissues is challenging, often resulting in redundant efforts.

Recognizing the limitations, a number of supervised approaches were proposed to utilize existing reference datasets as training dataset and directly annotate cells on query datasets without the clustering step. Mainstream numerical methods in this category include Seurat^9^ and SingleR^10^. Seurat identifies anchor across batches using mutual nearest neighbour and then use supervised PCA on mutual neighbours to transfer reference annotations^9^. Meanwhile, SingleR utilize the Spearman correlation-based scoring to perform annotation transfer task^10^. However, both of these methods have poor scalability that led to longer runtimes and higher memory usage^11^.

Deep learning methods offer a promising solution for supervised label transfer, where one popular approach involves training a neural network to perform representation learning. Representation learning is the process of automatically extracting meaningful features or representations from raw data, aiming to create a more concise and informative representation that captures the underlying patterns and structure. These learned representations can then be utilized for downstream annotation transfer tasks. In the realm of representation learning models, scANVI^12^ and Concerto^13^ are among the state-of-the-art approaches. The scANVI model employs variational inference deep generative model to learn a compact representation of gene expression patterns in single-cell RNA sequencing (scRNA-seq) data. It integrates this learned representation with a reference dataset to transfer annotations from the reference to the query dataset. On the other hand, Concerto utilizes a contrastive learning approach to learn a low-dimensional embedding. It leverages K-nearest neighbors (KNN) in this embedding space to predict annotations for the query dataset. Both methods have demonstrated remarkable performance in label transfer tasks, surpassing other existing methods^11–13^.

In this study, we introduce a novel framework named Supervised Contrastive Learning for Single Cell (SCLSC) for single-cell type annotation. SCLSC leverages supervised contrastive learning, which utilizes label information from the training data to provide explicit guidance on the similarity or dissimilarity between samples during the learning process. We devised supervised contrastive loss that use cell type representations. These cell type representations can guide the model to position a sample in proximity to its respective representative cell within the embedding space while ensuring it remains distant from representative cells belonging to different labels. Through a comprehensive evaluation using both real and simulated datasets, we demonstrate that the learned representations from SCLSC offer several advantages. First, they enable improved separation of cell types, effectively handling variations and noise present in the data. Second, using cell type representation simplify the contrastive learning process because the number of cell type is inherently less than the number of cells. It make SCLSC a fast and straightforward framework that scales well, even when dealing with large datasets. These characteristics make SCLSC particularly suitable for practical applications where computational efficiency is essential. Additionally, the representations capture relevant information for the label transfer task, leading to enhanced performance compared to existing state-of-the-art methods. Overall, our findings highlight the effectiveness of the supervised contrastive learning approach employed in SCLSC, showcasing its ability to achieve superior performance in single-cell label transfer tasks while maintaining simplicity, scalability, and efficiency.

## Results

### Overview of the SCLSC pipeline

We first introduced the complete pipeline of SCLSC. The SCLSC pipeline is divided into two main components: (1) embedding learning for cell and cell type and (2) cell annotation, shown in Figure 1. For representing scRNA-seq data, in previous work, highly variable genes across different cell types are used as the cell profiles. However, straightforward use for cell-type annotation is less comprehensive and consistent due to data noise and batch effect. With this in mind, we design a method to map the raw gene profiles of each cell into a new embedding space. To learn embeddings of cell and cell types, we used an MLP (Multi-Layer Perceptron) encoder to translate a raw cell profile into a new embedding space that factors in its cell type annotation. Supervised contrastive learning is used to train the MLP encoder, allowing us to acquire such a new representation. To represent cell types in the same embedding space as single cells, we use the arithmetic mean of gene profile vectors from all cells annotated with the cell type as an approximation. The model parameters of the MLP encoder are shared between both cell and cell types. We optimized the supervised contrastive loss between the cell samples and the cell type representative for updating the MLP encoder. Through the supervised contrastive learning process, cells of the same type tend to become more clustered, while cells from different cell types are increasingly separate. As a result, the cell type annotation is transferred through the embedding mapping for other scRNA-seq profiles. Once the MLP encoder is learned, a KNN is then trained within the new embedding space. Subsequently, a new cell can be annotated based on the KNN within this new embedding space, as depicted in Figure 1b.

**Figure 1.**
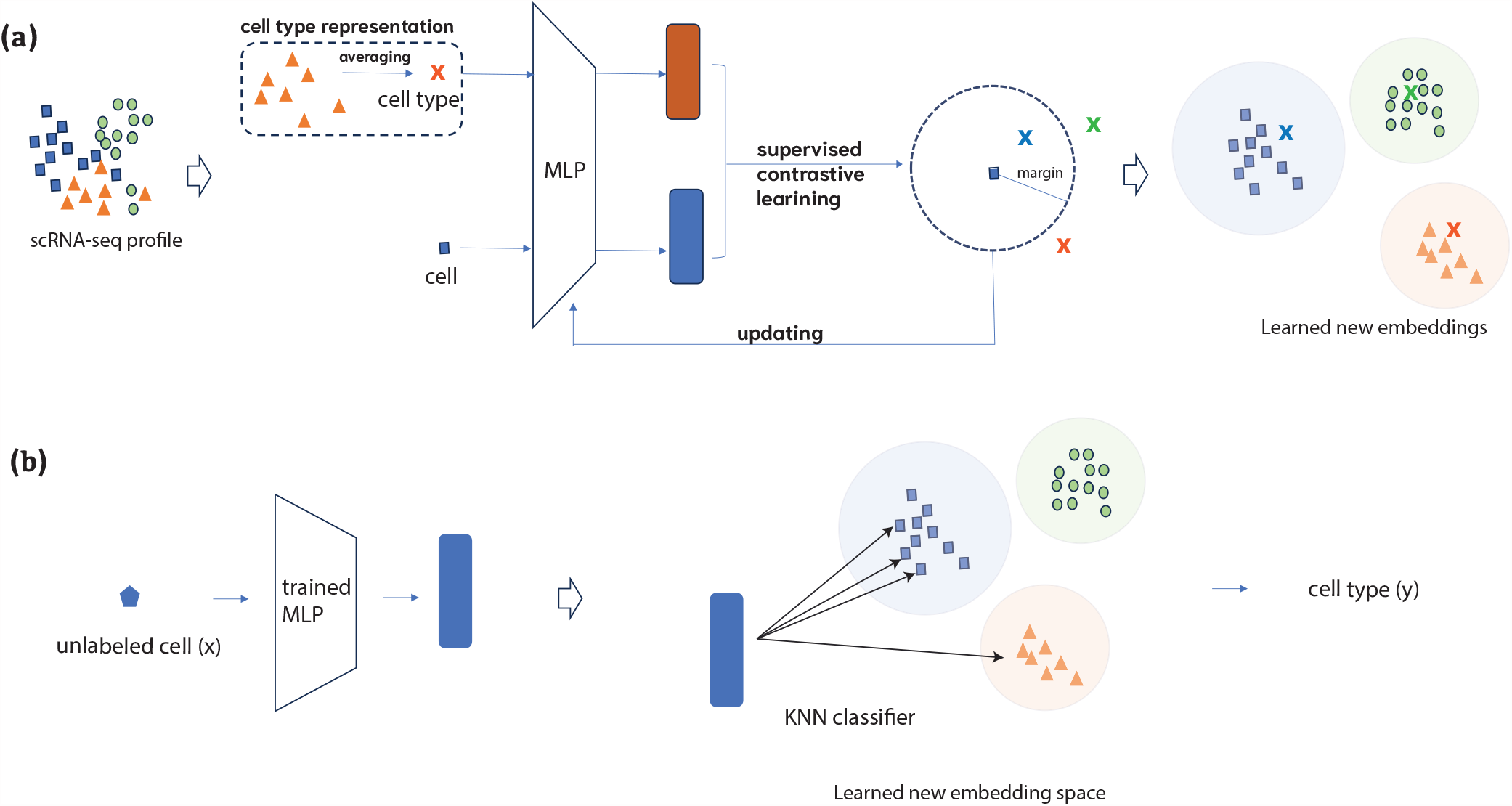
Overall view of SCLSC pipeline. The SCLSC pipeline can be divided into two phases: embedding learning and cell type annotation. In the first stage, as shown in (a), supervised contrastive learning is applied to learn new embeddings that capture the cell and cell type relationship derived from supervised data. For cell type representation, we averaged cell profile vectors in the same cluster as an approximated cell type profiling. In the second stage, as shown in (b), a candidate cell profile is mapped to its new embedding space based on the learned encoder in the first stage. Then, KNN is applied in the new embedding space to assign the cell type annotation for the cell.

### SCLSC achieved state-of-the-art label transfer task performance

Here, we evaluate the performance of SCLSC for label transfer task. The label transfer task refers to the process of transferring known labels or annotations from one dataset to another. Our approach involves several steps: (1) calculating query embeddings using pretrained model weights, (2) locate query cells near their most similar reference cells, and (3) use a KNN classifier (with a default value of k = 10) to transfer reference annotations to the query cells.

To evaluate the effectiveness of SCLSC, we compare its performance against other methods, including Seurat based on mutual nearest neighbors, SingleR based on correlation, scANVI based on variational inference probabilistic model, and Concerto based on contrastive learning. Two experiments were designed for evaluation: the random split experiment and the split by batch experiment. In the random split experiments, we divided the dataset into training, validation, and test datasets using stratified random splits. On the other hand, in the split by batch experiment, we divided the dataset based on the batches to assess the performance under batch-specific conditions.

Based on our experiment with random splits, it was consistently observed that SCLSC consistently achieves the highest accuracy and macro-averaged F1 score across most datasets, except for the lung and pancreas dataset where it comes in second place. (Figure 2a-2b). Particularly in the PBMC dataset, where it achieved 12% and 27% improvements of accuracy and macro-F1 score, respectively, compared to the second-ranked method. It is worth noting that each dataset has its own distinct characteristics. In the case of the PBMC dataset, challenges in cell type classification arose due to imbalanced distributions among cell types and strong correlations between them. Notably, the correlation-based method SingleR exhibited the poorest performance in this dataset, likely due to the presence of multicollinearity issues. Moving on to the CeNGEN C. Elegans dataset, it is characterized by a large number of cell types, with some cell types being rare, consisting of less than 100 cells. The neural network approach utilizing a variational inference probabilistic model, scANVI, demonstrated subpar performance in this dataset. In the case of the Thymus dataset, which contains a large number of samples, SingleR failed to produce any outputs even after running for more than 6 hours, prompting us to terminate the process. In contrast, the consistently superior accuracy of SCLSC across all datasets demonstrates its efficacy in addressing challenges such as multicollinearity problems, imbalanced distribution of cell types, and large-scale samples. Additionally, SCLSC consistently achieves the highest macro-averaged F1 score across most of the datasets. The macro-averaged F1 score serves as a valuable performance metric, particularly in scenarios where class imbalance exists. It calculates the F1 score for each class individually and computes the average of these scores. This approach ensures that each class is given equal importance, irrespective of its prevalence in the dataset. Based on the high macro-averaged F1 score, SCLSC demonstrates superior ability in accurately separating rare cell types (Figure 2a, Figure S1).

**Figure 2.**
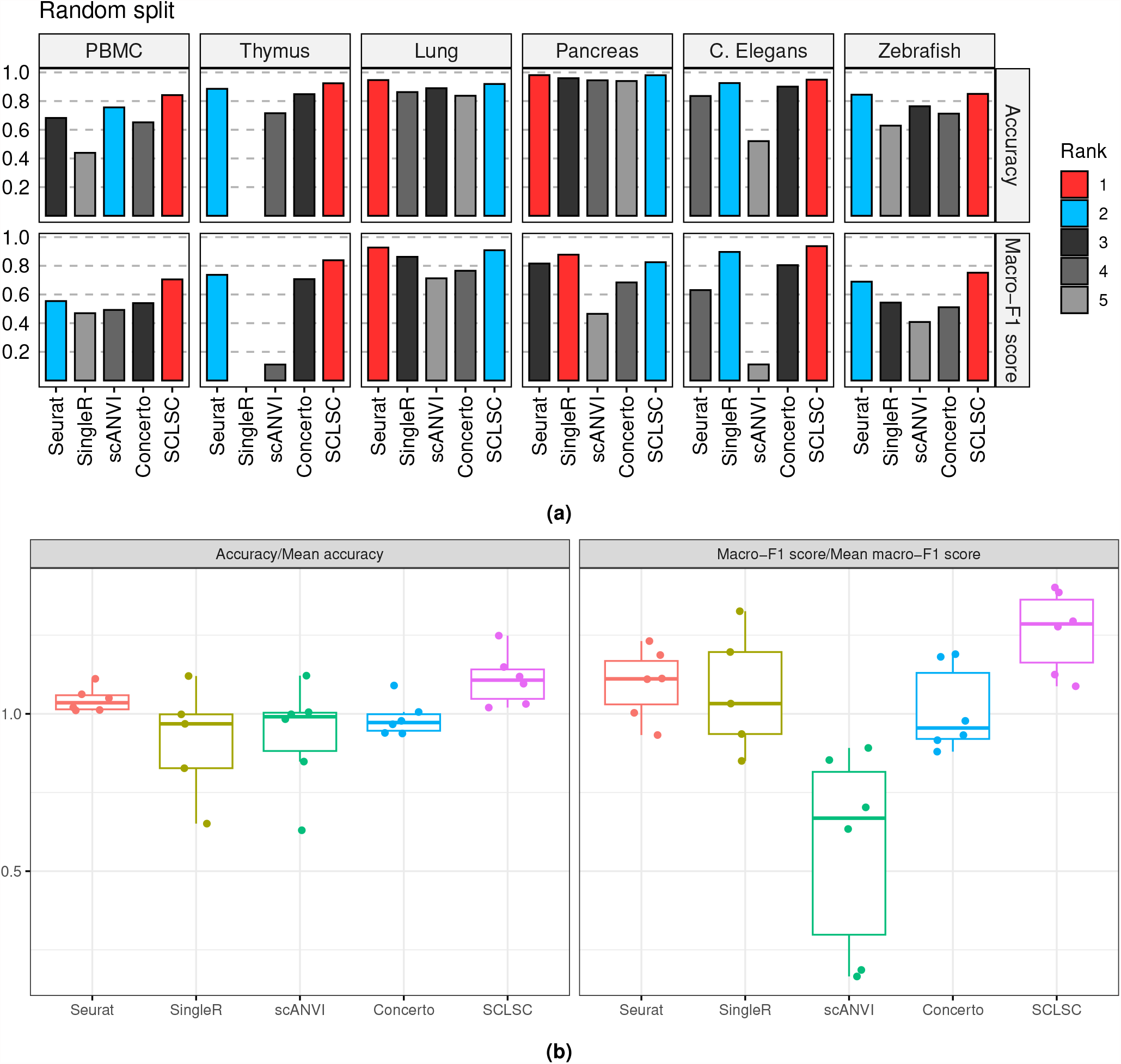
SCLSC achieves superior accuracy for label projection tasks. (a) The performance of SCLSC in the label transfer task surpassed that of other state-of-the-art methods, as evidenced by its high accuracy and macro-averaged F1 score across six benchmark datasets.(b) SCLSC demonstrated performance above average when comparing its relative accuracy to the mean accuracy and the relative macro-F1 to the mean macro-F1 score across different datasets.

### Cell type hierarchy is preserved in the new embedding space

In conducting supervised contrastive learning, we approximated the cell type representation by using the arithmetic mean of gene profile vectors from all cells annotated with a specific cell type. We utilized the PBMC dataset to investigate the cell type hierarchy for this method in both the raw and learned embedding spaces. Hierarchical agglomerative clustering with single-linkage was performed on 11 cell type vectors in each of these spaces. The dendogram of cell type representation from the 2000 HVGs input (Figure 3a) and the dendogram of the output cell type representation learned by SCLSC (Figure 3b) is consistent with the immune cell lineage tree (Figure 3c). The fact that the input HVGs dendrogram aligns with the immune cell differentiation hierarchy indicates that the selected input adequately captures the biological information within the dataset.

**Figure 3.**
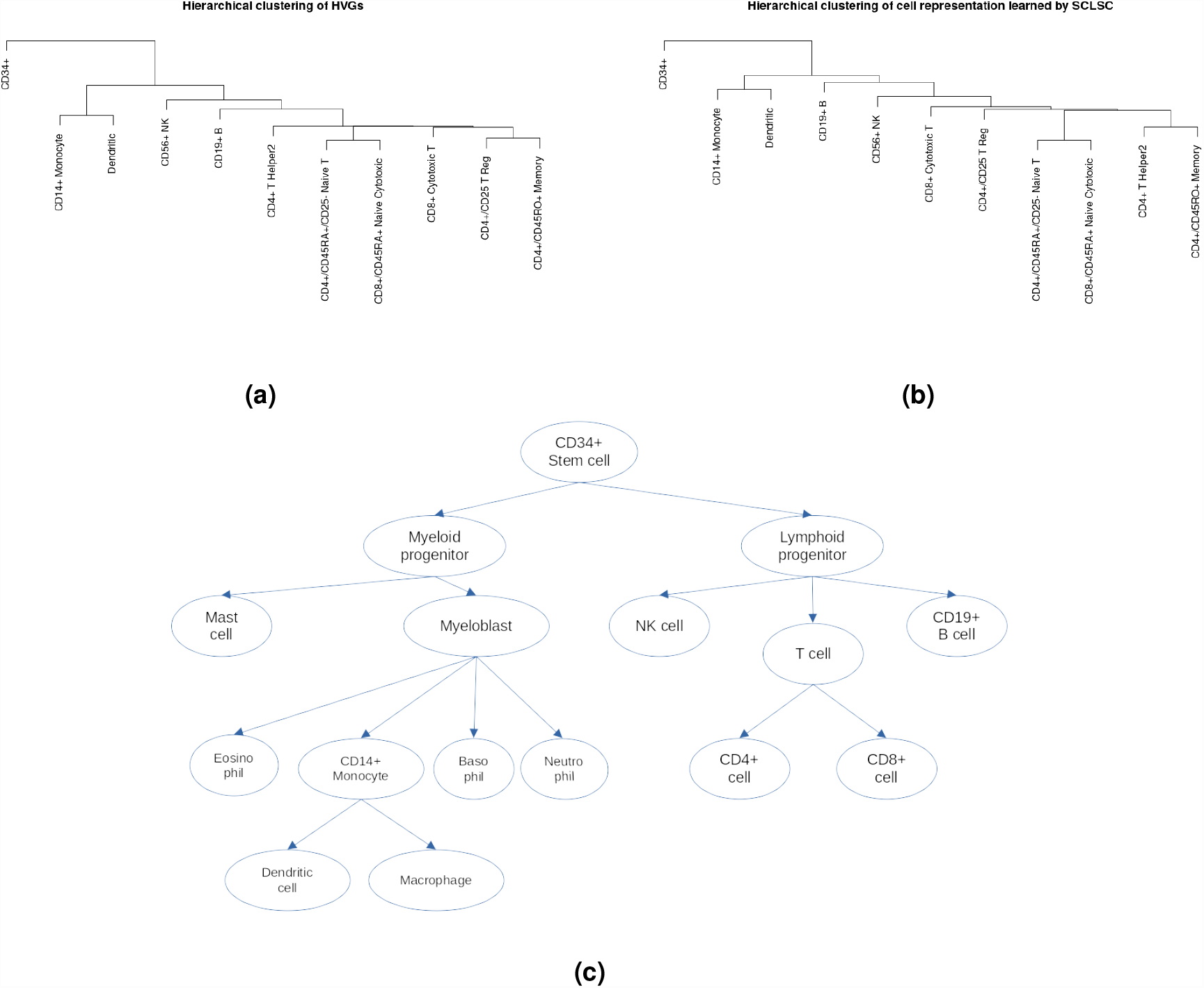
The comparison of dendrogram of PBMC cell type representation and the hierarchy of immune cells differentiation.(a) The dendrogram of the cell type representation using HVGs in original space that being used as input and **(b)** the dendrogram of the cell type representation learned by SCLSC showed that SLCSC input and output can capture and preserve **(c)** the hierarchy of immune cells differentiation.

In the dendrogram shown in Figure 3b, CD34+ cells are positioned at the root, from which two branches emerge. The left branch consists of myeloid cells, including CD14+ monocytes and dendritic cells, while the right branch consists of lymphoid lineage cells, namely CD56+ NK, CD19+ B, CD4+ T, and CD8+ T cells. This dendrogram structure aligns with the established immune cell lineage tree, demonstrating that the cell type representation learned by the SCLSC effectively captures and preserves the biological hierarchical structure present in the dataset.

### The effectiveness of SCLSC for label transfer task under batch specific conditions

To investigate the performance under batch-specific conditions, we also divided some datasets based on their batches. Although the SCLSC method does not directly tackle the batch effect issue, our findings demonstrate that SCLSC performs well even in the presence of batch effects (Figure 4a). SCLSC has the high accuracy and macro averaged F1 scores, ranked first or second, in all dataset. There are two plausible explanations as to why our model can partially address the batch effect problem. First, contrastive learning commonly employs data augmentation techniques that introduce diverse transformations to the input data. In this context, the batch condition can be viewed as a form of data augmentation^14^. By leveraging contrastive learning, our model learns to emphasize the shared features among samples while remaining invariant to batch-specific variations. Consequently, this diminishes the impact of batch effects on the learned representations. Second, it is widely accepted that the differences between cell types are greater than differences between batch^15,16^. As SCLSC functions as a supervised model, it utilizes cell type information to guide the learning process and minimize the influence of batch effects.

**Figure 4.**
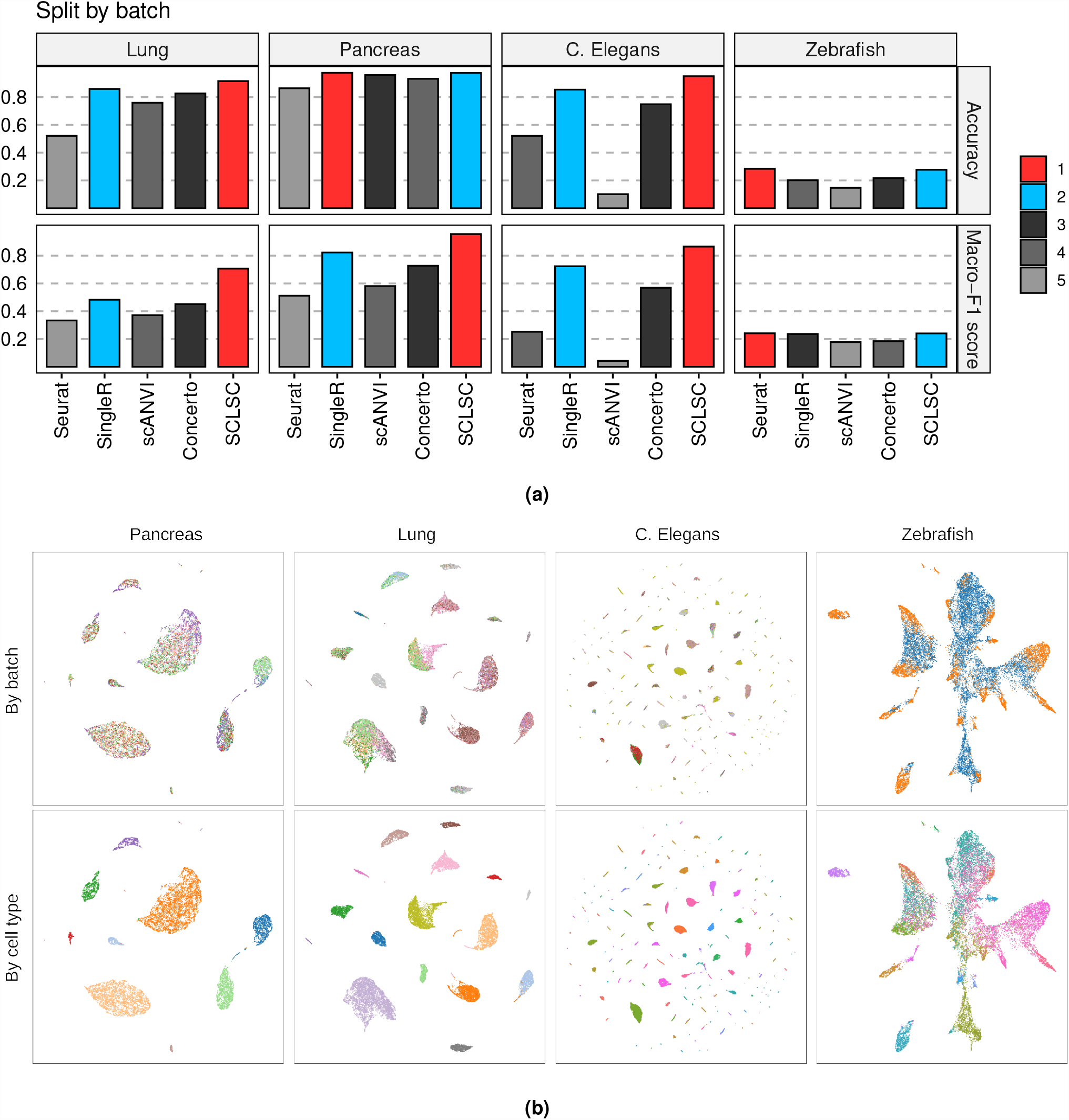
SCLSC can handle label transfer task under batch effects. (a)The incorporation of label information in SCLSC can tackles the batch effect, as demonstrated by its high accuracy and macro-F1 scores across benchmark datasets. (b)The benchmark datasets’ learned representation by SCLSC is visualized using UMAP. The upper figure represents the color-coding of batches, while the lower figure represents the color-coding of cell types. The batch groups are well mixed within each cluster, while simultaneously maintaining clear separation of individual cell types.

### SCLSC is fast and scale-well in large dataset

We used simulated datasets generated using Splatter R package^17^ for scalability analysis. The dataset has 2000 features/genes and 10 cell types with with equal occurrence frequencies. Figure 5 showed that SCLSC could easily deal with large-scale datasets. The SCLSC demonstrates an almost twofold increase in speed compared to the second fastest techniques, Concerto, when applied to a dataset consisting of 400,000 samples. SCLSC speed stems from a combination of the simplicity of the contrastive learning algorithm, the straightforwardness of the encoder architecture, and the support for GPU acceleration. The Early Stopping algorithm integrated into the SCLSC also has role to stop the training process early if the model’s performance is not improving, thereby saving time and computational resources. The SingleR method was excluded from the scalability analysis alongside other methods due to its failure to generate results within the given time frame for thymus datasets.

**Figure 5.**
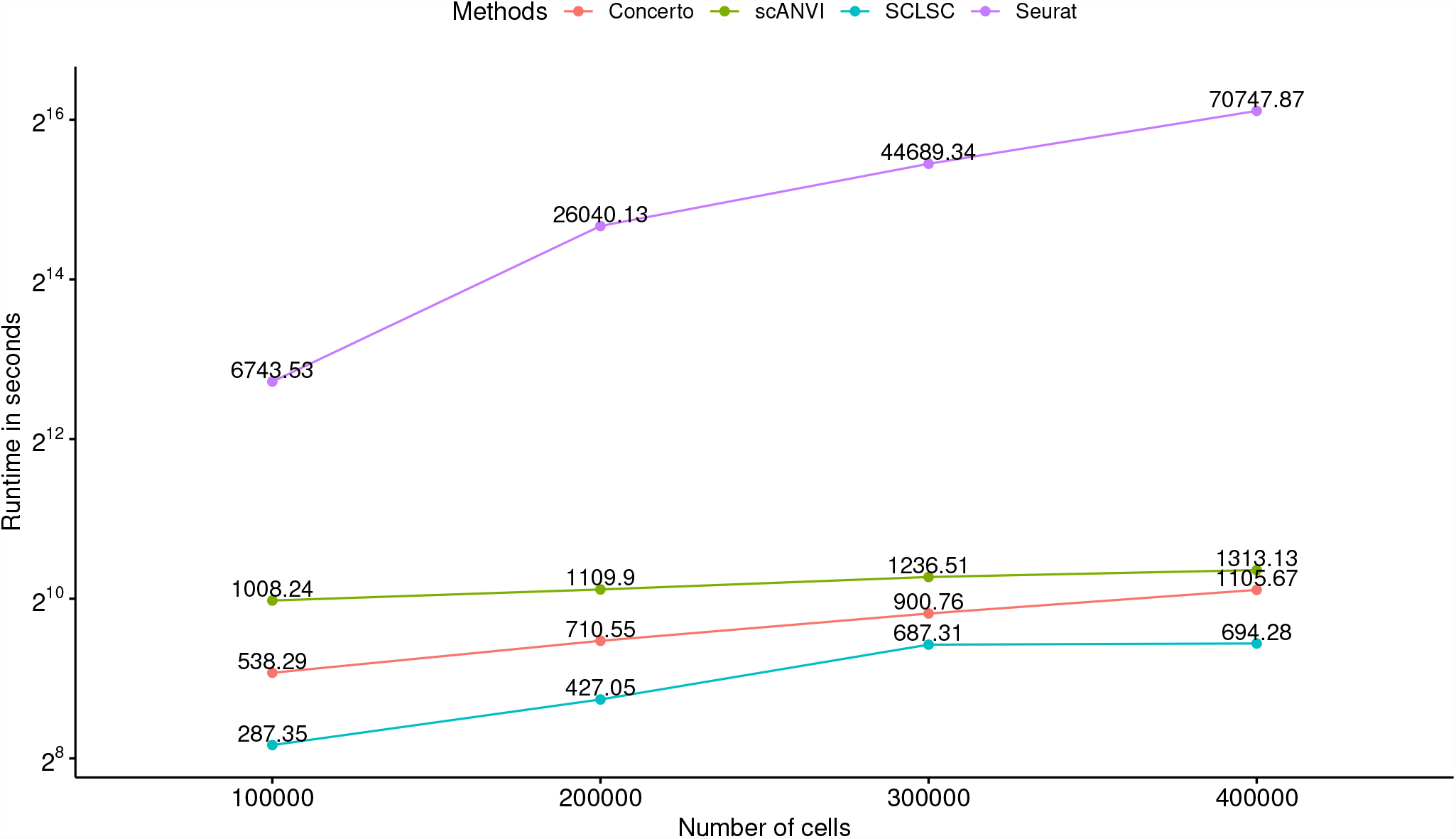
The runtime of the SCLSC and the competing methods against the number of cells. Due to its simplicity, the SCLSC algorithm is capable of efficiently scaling up to handle large datasets. Our findings demonstrate that, when applied to query-to-reference label transfer tasks involving 100,000 to 400,000 cells, the SCLSC algorithm outperforms other methods in terms of speed, even almost reaching two times faster than second rank methods in analyzing 400,000 cells.

### Real world application: mapping label from PBMC dataset onto immune cells from dengue datasets

As an example of real-world implementation, we use SCLSC to project cell labels from PBMC dataset from Zheng et al.^18^ onto PBMC from a patient with dengue fever (DF) and a patient with dengue hemorrhagic fever (DHF)^19^. The PBMC samples of dengue dataset were gathered on specific days: at defervescence (Def) day, two days before defervescence (Day-2), one day before defervescence (Day-1) - collectively known as the febrile illness period, and two weeks after defervescence (Wk2), which represents the convalescence or follow-up phase. SCLSC can successfully transfer the label from PBMC dataset onto unlabelled cells in the dengue dataset (Figure 6a). With the aid of these labelled cells, we can conduct downstream analysis to track the dynamics of immune cell populations during dengue virus infection, focusing on B cells in particular. Recent observations have emphasized the significant involvement of B cells during infection with dengue viruses, particularly during acute dengue infection, where a substantial increase in the number of effector B cells has been noticed^20^. To verify the accuracy of the labelling process, we conducted differential gene analysis and gene set enrichment analysis on cells identified as CD19+ B cells. The results confirmed that the labelled cells accurately represented the genetic characteristics associated with B cell cellular processes, as they exhibited enrichment of relevant genes (Figure 6b). As shown in the Figure 6c, the proportion of the B cell is peaking in the one day before defervescence. This finding is consistent with a previous study that showed an increase in immunoglobulin-containing B cells can be observed during infection, and these cells reach their maximum levels around the time when the fever starts to subside^21,22^. From the cluster of B cells, we can also investigate the subtype of the B cells suc as plasmablast and plasma cell. For this purpose, we used gene markers, namely XBP1^23^, TNFRSF17^24^, and JCHAIN^25^, which are plasma cells and plasmablasts markers. On the other hand, we used MS4A1^26^ as a gene marker for B cells, excluding plasmablasts and plasma cells. As shown in the Figure 6d, the SCLSC learned the representation that clearly separate the effector B cells (plasmablast and plasma cell) and the non-effector B cells.

**Figure 6.**
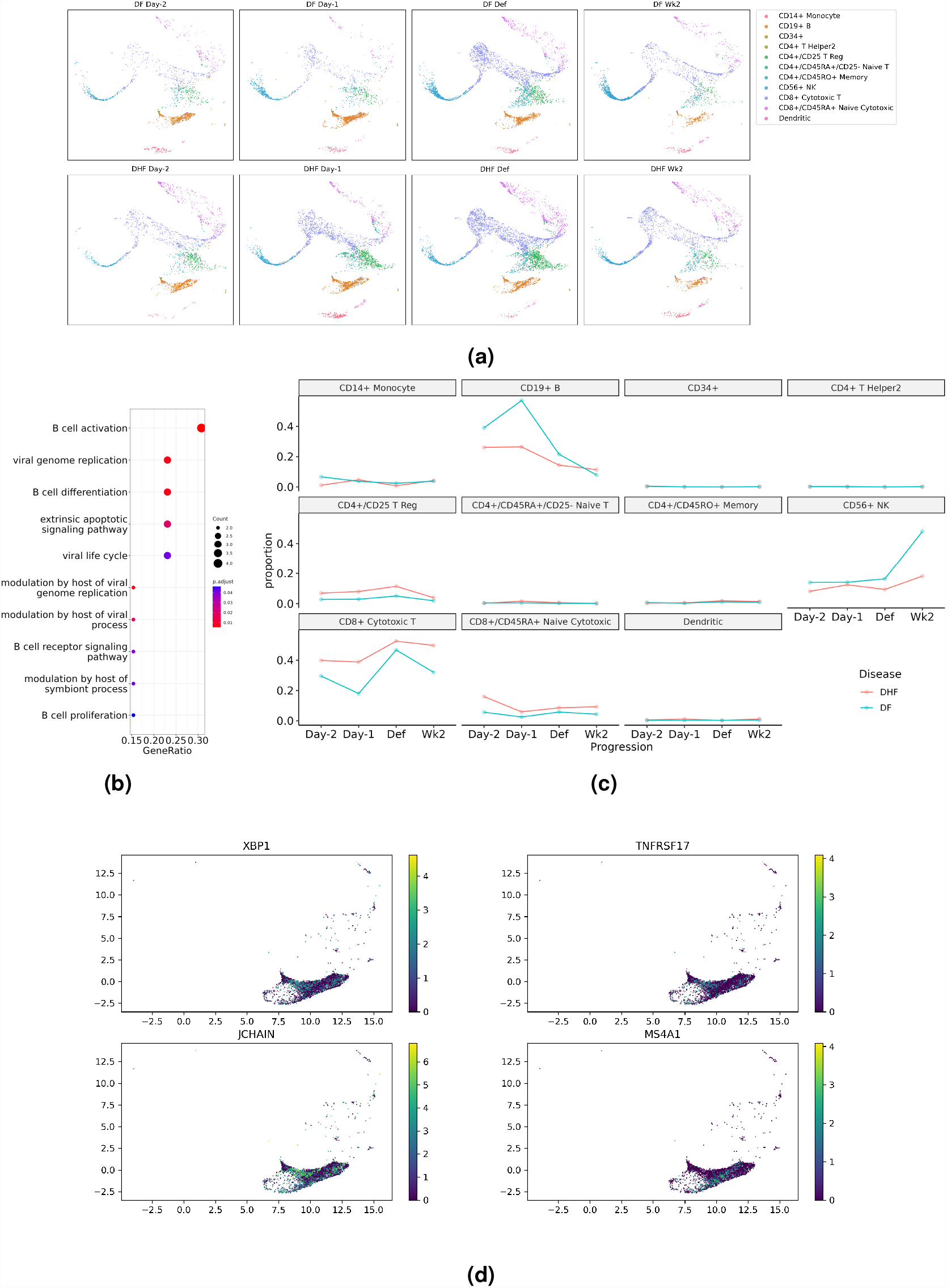
Downstream analysis of the representation of PBMC dengue dataset learned by SCLSC (a) UMAP visualization of the representation of PBMC dataset learned by SCLSC. **(b)** The presence of enriched genes associated with B cell cellular processes in the cells labeled as CD19+ B confirms the accuracy of our labeling, confirming that these cells are indeed B cells. **(c)** The predicted cell labels can be utilized for subsequent downstream analysis, including monitoring the dynamic changes in subpopulations of PBMC cells over dengue infection progression. **d** The learned representation of B cells using SCLSC demonstrates the ability to distinguish subtypes that were not observed in the training dataset. The representation separates the plasmablast/plasma cell subtype, characterized by XBP1, TNFRSF17, and JCHAIN markers, from the less differentiated B cells labeled by MS4A1.

### SCLSC provides stable performance across varying dimensions of input and output

SCLSC has two key parameters: the dimension of the input and the dimension of the output of the encoder. In case of input dimension, SCLSC has the capability to process input from all genes. However, a subset of genes known as highly variable genes (HVGs), which exhibit high cell-to-cell variation, have been found can help in reducing noise and emphasizing the biological signal in scRNA-seq datasets^9,27^. To investigate this further, we conducted an experiment using both all genes and HVGs as SCLSC inputs for the label transfer task on benchmark datasets. We found that using all genes as inputs did not lead to a significant difference in accuracy (*n* = 5, Wilcoxon test *P*-value = 0.11) and made the macro F1-score worse (*n* = 5, Wilcoxon test *P*-value = 0.0311) compared to using HVGs as inputs (Figure 7a). Furthermore, utilizing all genes required more time and memory resources.

**Figure 7.**
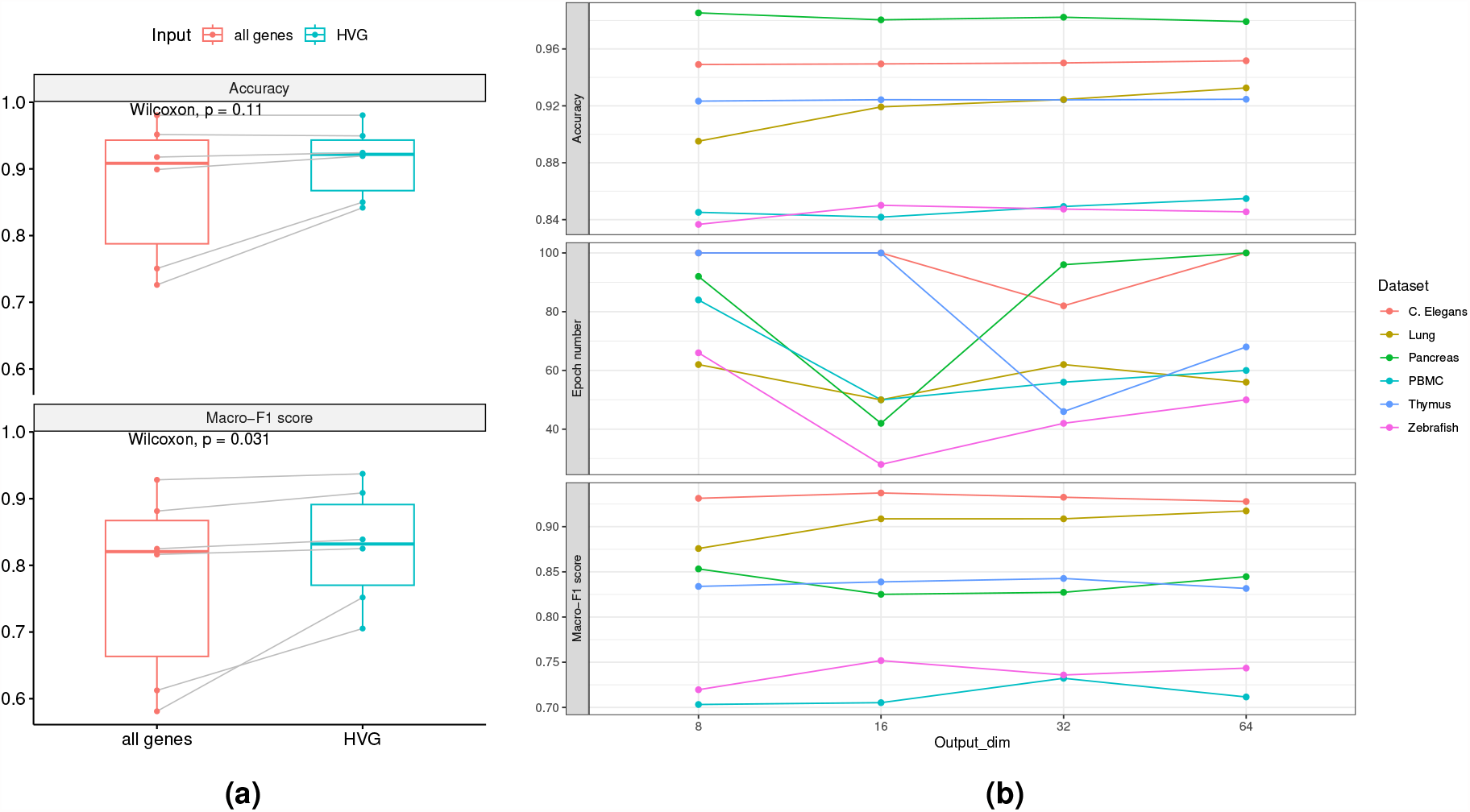
The impact of the encoders input and output dimension to the performance of the SCLSC (a) Using all genes as input does not improve accuracy (*n* = 5, Wilcoxon test *P*-value = 0.11), in fact, can be detrimental to the macro-F1 score of SCLSC (*n* = 5, Wilcoxon test *P*-value = 0.0311). **(b)** On the different choice of output dimensions, there were no significant changes in accuracy and macro-F1 score observed. However, the epoch needed to trigger the early stopper is decreased when the output dimension is 16 in most datasets.

We also conducted an experiment using encoder output dimension = 8,16,32, and 64. We observed that the tested output dimension had minimal impact on both the accuracy and macro-F1 score. However, the number of epochs needed until the early stopper was triggered decreased when the output dimension was set to 16 in the majority of the datasets (Figure 7b). This suggests that an output dimension of 16 can achieve optimal accuracy in the shortest time compared to other output dimensions. Based on these results, we used the 2000 HVGs as the input and 16 as the output dimension in our overall study.

## Discussion

### Methodology differences compared with previous contrastive learning method for single-cell data

In this research, we used real and simulated testing datasets to showcase the effectiveness of SCLSC in transfer cell type knowledge from annotated dataset. One key advantage of the SCLSC method is its simplicity for conducting contrastive learning for cell and cell types. Unlike the previous method doing contrastive learning for different cells, using cell types in the contrastive learning significantly reduced the number of contrastive pairs. With such a feature, SCLSC can tackle diverse datasets efficiently. SCLSC has capability to swiftly provide cell type information for hundred-thousands of cells in a matter of minutes. In our experiment, SCLSC proved to be the fastest, outperforming the second-best method by 37% in processing 400,000 cells.

Previous studies have shown that contrastive learning models possess a remarkable ability to transform single cell input data into a more compact, structured, and expressive representation that captures data characteristics^13,28^. However, by incorporating the label information, supervised contrastive learning have several advantages compared to conventional contrastive learning. First, the model can learn representations that are optimized specifically for the label transfer task, resulting in more distinguishable outcomes. Second, it reduced the sensitivity to negative samples. In conventional contrastive learning, the choice of negative samples plays a crucial role in the learning process. Selecting appropriate negative samples can be challenging, and suboptimal choices may result in the model learning trivial or uninformative patterns. Supervised contrastive learning, by incorporating label information, reduces the reliance on negative samples, making it more robust and less sensitive to the choice of negatives. Third, it is enabling more effective training even with smaller labeled datasets. In traditional contrastive learning approaches, a large labeled dataset is typically required to capture similarity and dissimilarity patterns effectively. However, supervised contrastive learning can leverage labeled data, enabling more effective training even with smaller labeled datasets. Moreover, SCLSC algorithm is using cell type representation, which are the average gene expression of the cell type, to perform cell-cell type pairs instead of cell-cell pairs for contrastive learning. The number of cell types is significantly smaller than the number of cells. Thus, performing contrastive learning is easier and faster because we just need to compare the cell with small number of cell types rather than thousands of cells. Compared to our method, Concerto^13^ utilizes unsupervised contrastive learning and employs an asymmetric teacher-student architecture to achieve high performance. In contrast, SCLSC employs a simpler MLP but still achieves comparable results. To assess the agreement between SCLSC and Concerto on the PBMC dataset, we calculated the Normalized Mutual Information (NMI) score, which turned out to be 0.48206. This score clearly indicates that our approach, supervised contrastive learning, differs from Concerto’s unsupervised contrastive learning.

There is no universal model that fits all scenarios, and our SCLSC model is no exception. The validity of our model heavily relies on the availability and accuracy of labels in the training data. Acquiring a large amount of precisely labeled single-cell data poses a challenge in the field of single-cell research. Therefore, the applicability of the SCLSC model is restricted to tasks where labeled data is readily accessible. Utilizing high-quality reference datasets leads to superior annotations and enhances the model’s performance. Additionally, the generalization capability of the model is limited when it comes to unseen classes. If the test or deployment data includes classes or samples that were not present during training, the representations produced by the SCLSC model may not generalize well to these unseen classes. To mitigate this limitation, we recommend using large-scale reference datasets that encompass a wide range of cell types during the training process. It’s important to consider these limitations when applying the SCLSC model in practical applications and to take appropriate measures such as addressing batch effects, ensuring sufficient availability of accurately labeled data, and using comprehensive reference datasets to enhance generalization to unseen classes.

### Data

#### Benchmark dataset

We have chosen commonly utilized datasets for benchmarking label transfer tasks. These datasets are publicly accessible and come in a standardized format that is ready for immediate use. Each dataset possesses distinct attributes, including imbalanced distributions among cell types, multicollinearity between cell types, a large number of cell types, a substantial number of cells, and challenges related to batch effects.

#### PBMC dataset

We used Zheng *et al*^18^ Peripheral Blood Mononuclear Cells (PBMC) dataset freely available from 10X Genomics. After preprocessing, we got total 68265 cells before splitting. This dataset contains rare cell types and the distribution of cell types is imbalanced. Moreover, the cell types in the dataset were highly correlated with each other make it difficult to differentiate them.

#### Pancreas dataset

The pancreas dataset is a human pancreatic islet scRNA-seq data from 6 sequencing technologies (CEL-seq, CEL-seq2, Smart-seq2, inDrop, Fluidigm C1, and SMARTER-seq)^11^. We splitted the dataset by technology batch (training: 8 batches, test: 1 batch) for batch effect investigation.

#### Lung dataset

Human lung scRNA-seq data were obtained from 2 different sequencing technologies: 10X and Drop-seq. We splitted the dataset by technology batch (training: 15 batches, test: 1 batch) for batch effect investigation^11^.

#### Thymus dataset

The thymus dataset were from the single-cell RNA sequencing of cells inside human thymus^29^. This large dataset contains more than 250,000 cells which make it useful for evaluating methods scalability.

#### Zebrafish dataset

This dataset contains the data of zebrafish embryos cells during the first day of development, with and without a knockout of chordin, an important developmental gene. After preprocessing, this dataset had dimension 26022 cells × 2000 genes and 24 cell types.^30^. We splitted the dataset by laboratory for batch effect investigation.

#### CeNGEN dataset

This dataset are from The Complete Gene Expression Map of the C. elegans Nervous System (CeNGEN) project and contains FACS-isolated C. elegans neurons data sequenced on 10x Genomics^31^. This dataset is characterized by a large number of cell types and imbalanced distribution of cell types. We splitted the dataset by batch (training: 12 batches, test: 5 batches) for batch effect investigation.

#### Dengue dataset

These data originate from PBMC cells taken from patients with acute dengue virus infection^19^. Because this dataset lacks labels, we utilized a PBMC dataset as a reference to annotate the cells in the dengue dataset for real-world application experiments.

### Methods

#### Training process

We trained the MLP using gene expression count matrix and representative samples as the inputs. A representative sample corresponding to a label was defined as the mean of the all samples that shared the same label. After selected the representative samples, we embed the gene expression matrix and representative samples using MLP encoder. In embedding space, we calculated the supervised contrastive loss between the samples and the representative samples for updating the MLP encoder. To mitigate the possibility of the overfitting, we employed an early stopping algorithm. In this algorithm, the training will stop if the validation loss is not decreasing for five validation steps. A validation step was performed for every two epoch training.

### SCLSC Supervised Contrastive loss

We optimized the contrastive loss, defining it as follows:

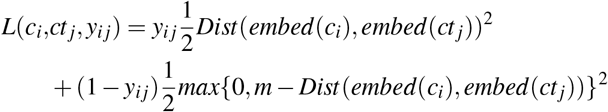

The variables *c*_*i*_, *ct* _*j*_, *y*_*ij*_ are the cell profiles, the cell type representation profiles, and the one-hot-encoded cell label, respectively. The cell type *ct* _*j*_ is computed using all cells of that type from the training data. It is calculating as 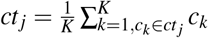. Here, K represents the number of cell samples of the type *ct* _*j*_. A margin *m* is employed to group cells within their respective annotated cell type, while distinguishing them from the other cell types.

### Encoder structure

The encoder network accepts *X* ∈ ℝ^*d*^ where *d* denotes the number of genes. First, the encoder feed X into a dense layer with Relu activation, dropout, and batch normalization layer to get **hidden**_**1**_. Then, the **hidden**_**1**_ is fed into a second dense layer with Relu activation, dropout, and batch normalization layer to get **hidden**_**2**_. Finally, the **hidden**_**2**_ is fed into a last dense layer to get the final output *Z* ∈ ℝ^*d*^′ where *d*^′^ denotes the number of embedding space dimensions.

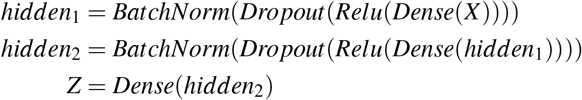

### Compared baseline methods

#### Seurat

Seurat is a widely used R package that has been developed for analysis and exploration of single-cell RNA sequencing data. Seurat transfer the annotation data by using a canonical correlation analysis of a set of anchor genes that are highly variable and shared between the reference and new datasets^9^. We used Seurat v4.0 and followed the transfer annotation from query datasets tutorial to perform label projection in this study.

#### SingleR

SingleR is an automatic annotation method for single-cell RNA sequencing implemented in R package^10^. In the SingleR pipeline, a Spearman coefficient is calculated for single-cell gene expression with each of the samples in the reference dataset, using only the variable genes in the reference dataset to increase the ability to distinguish closely related cell types. This process is performed iteratively using only the top cell types from the previous step and the variable genes among them until only one cell type remains. We used default parameter of SingleR R package in our study.

#### scANVI

scANVI is a semi-supervised model that employed Variational Inference to annotate a dataset of unlabelled query dataset from annotated reference dataset^12^. In our study, we used scANVI methods implemented in scvi-tools Python package with default parameter. To perform label projection, we followed the reference mapping scvi-tools tutorial.

#### Concerto

Concerto leverages a self-distillation contrastive learning framework to learn cell embeddings in lower dimensional space^13^. The learned cell embeddings are fed into KNN classifier to perform label projection task. In our study, we employed the Concerto source code downloaded from public repository (https://github.com/melobio/Concerto-reproducibility. We used default parameter stated in the Concerto source code.

### Data preprocessing

The datasets underwent preprocessing to eliminate cells with high mitochondrial gene expression (more than 5 percents of the cell total count), cells with minimal gene expression (number of genes per cell < 200), and genes that were only detected in a small number of cells (number of cells that expressed the gene < 3). Subsequently, We selected 2000 highly variable genes (HGV) using analytic Pearson residuals implemented in Scanpy package. Following this, we normalized the count of each cell to 10,000 counts and applied a *log*(*x* + 1) transformation. The resulting dataset was then divided into training, validation, and test sets with a ratio of 8 : 1 : 1. All of the preprocessing steps were performed using Scanpy package^32^. The summary of the dataset, reference, and download link were provided in Table 1.

**Table 1.** The summary of the datasets used in our study.

| Name | Description | No. cells | No. types | Download link | Ref |
| --- | --- | --- | --- | --- | --- |
| PBMC | Human PBMC cells | 68265 | 11 | PBMC | 18 |
| Pancreas | Human pancreas cells | 16382 | 14 | Pancreas | 11 |
| Thymus | Human thymus cells | 223792 | 44 | Thymus link | 29 |
| Lung | Human lung cells | 30717 | 17 | Lung | 11 |
| CeNGEN | C. Elegans neuron cells | 72857 | 169 | CeNGEN | 31 |
| Zebrafish | Zebrafish embryo cells | 26022 | 24 | zebrafish | 30 |
| Dengue dataset | PBMC of dengue fever patients | 39591 | NA | Dengue | 19 |
| Simulated dataset | Simulated dataset created by Splatter package | 500000 | 10 | - | 17 |

### Scalability analysis

For scalability analysis, simulated datasets were generated using Splatter R package^17^. The simulated datasets were generated using the following parameters: nGenes=2000 and group.prob=rep(1/10,times=10), while leaving the other parameters at their default values. The number of cells in the training dataset = 100000, 200000, 300000, and 400000 cells. The validation dataset and test dataset comprised 50,000 cells each, both derived from the same cell distribution as the training dataset. Then, each method was trained on each of the training datasets to make predictions regarding the annotations within the test dataset.

### UMAP visualization

Cells embedding were visualized by UMAP using umap-learn Python package. The umap parameters were set at their default values

### Dendogram visualization

First, we computed the averages of the 2000 High Variable Genes (HVGs) and the embedded features for each corresponding cell type, which yielded cell type representations in both the original input space and the embedding space. Then, we calculated the Euclidean distance matrix for these cell type representations in both spaces. Subsequently, we constructed dendrograms using single linkage based on the distance matrix. The dendograms were produced using hclust package.

### Dengue dataset analysis

We compute a ranking for the highly differential genes in the cells labelled as CD19+ B cells using Wilcoxon method implemented in the rank_genes_groups function in Scanpy package^32^. Next, we performed gene set enrichment analysis of 15 highest rank differential genes. The enrichment analysis were performed using clusterProfiler package^33^.

## Author contributions statement

Y.D.H. was responsible for the the data curation, analyses, and visualization, and writing the original draft of the manuscript. Y.Z. was responsible for the study conceptualization, data analysis, supervision, and editing the manuscript. S.I. was responsible for the funding acquisition, project administration, supervision, and editing the manuscript. All authors reviewed the manuscript.

## Data availability

All the real single-cell RNA sequencing datasets used in this study had been previously published. References to these datasets, information about their accessibility, and downloadable links can be found in Table 1.

## Code availability

The source code is accessible at https://github.com/yaozhong/SCLSC.

## Supplementary Material

**Figure S1.**
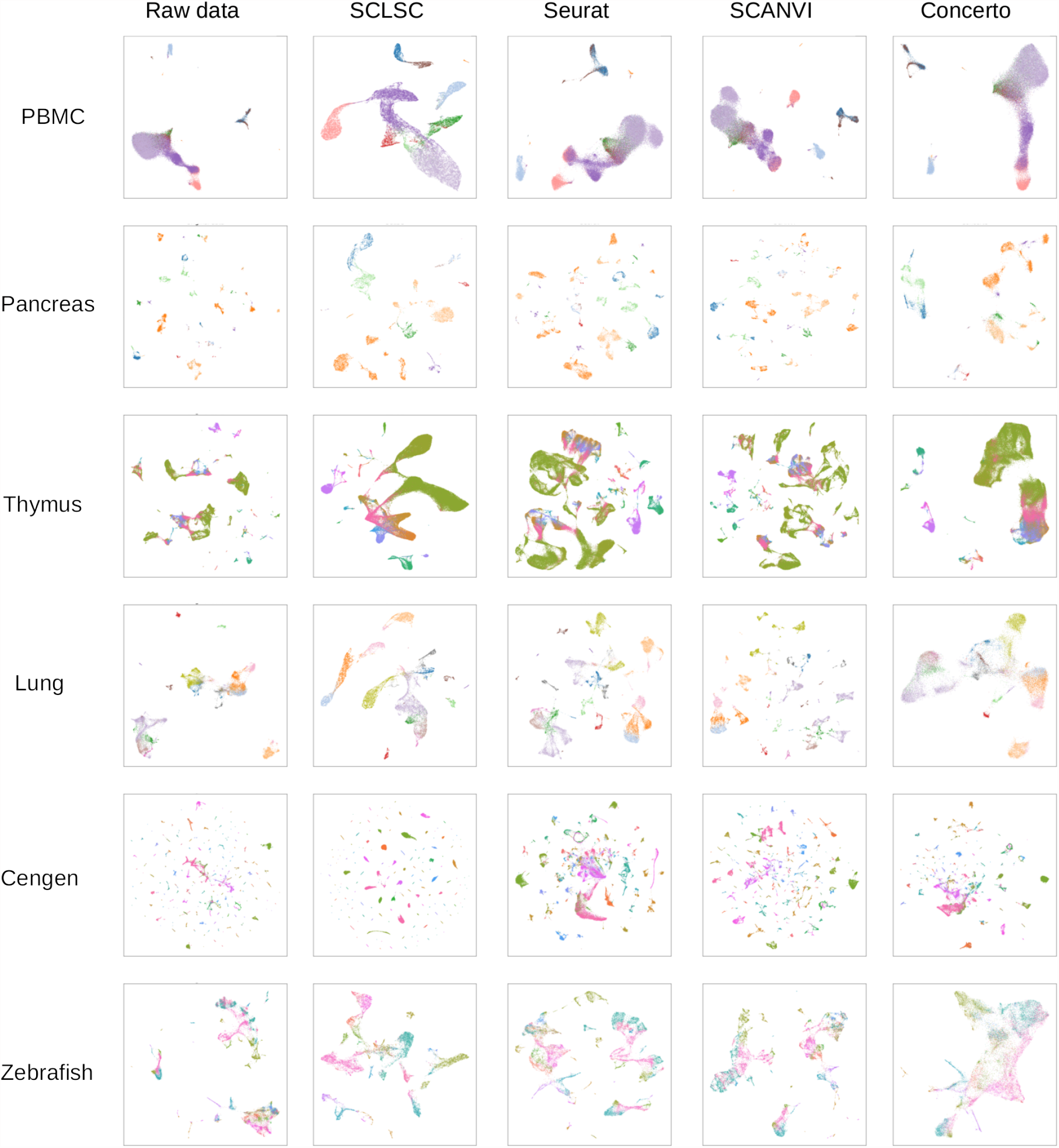
The UMAP visualization of benchmark datasets. SCLSC utilizes cell type representations to guide cells towards proximity to their respective cell type representations while simultaneously moving away from representations of other cell types. This process encourages cells of the same type to cluster together and separate from cells of different types, thereby facilitating more straightforward cell label predictions as shown in the visualization.

